# *Momordica charantia* Prevents Reproductive and Neurological Dysfunctions in *Drosophila melanogaster* Models of Type 2 Diabetes

**DOI:** 10.1101/2025.08.13.670173

**Authors:** A.M. Agi, R. Abdulazeez, N. Abdulasalam, M.B. Omobalaji, O.S. Haruna, D.M. Shehu

## Abstract

Type 2 diabetes (T2DM) poses a significant global health threat, compounded by factors such as obesity and lifestyle. This study investigates the anti-diabetic, reproductive, and neuroprotective effects of *Momordica charantia* (MC) in preventing T2DM. We used a high-calorie sucrose diet (2.5 g/10 g diet) to induce T2DM in Harwich and Indigenous strains of *Drosophila melanogaster*. Concurrently, we administered varying concentrations of MC (100 mg, 150 mg, and 200 mg/10 g diet) and 16 mg of metformin for 10 days. *M. charantia* significantly improved egg-laying ability in Indigenous and Harwich strains. Specifically, 150 mg and 200 mg doses significantly (p<0.01) enhanced egg-laying ability in Indigenous and Harwich strains, respectively. Additionally, 100 mg and 200 mg doses significantly rescued filial generation output in Harwich and Indigenous strains, respectively. *M. charantia* at the lowest dose (100 mg) reduced neuromuscular activity (p < 0.01) in both strains. Our results demonstrate that *M. charantia* prevents the onset of T2DM, improving reproductive fitness and reducing neuromuscular impairments associated with hyperglycemia. These findings suggest that *M. charantia* may be a promising therapeutic agent for preventing T2DM and its associated complications.

## Introduction

Type-2 diabetes (T2DM) is a combination of both genetic, environmental factors and lifestyle such as physical inactivity, obesity as a result imbalance diets (Ashcroft and Rorsman, 2018; Ismail *et al*., 2021). The prevalence of diabetes mellitus (DM) is currently estimated at 537 million surpassing the previously 400 million estimate and, is projected to increase to 783 million by 2045 of which 90 % have type-2 (Sun *et al*., 2022; WHO, 2022). In Africa, the incidence of DM is currently estimated to be about 24 million cases, which is projected to reach 55 million by 2045 (Sun *et al*., 2022). About 2 % of the 3.7 % prevalence of the more than 3.6 million affected by DM in Nigeria are found in Zaria province of Kaduna State (Dahiru *et al*., 2008; Wu *et al*., 2022).

*D. melanogaster*, known colloquially as the fruit fly, is common pest that is found in the environment such as homes, restaurants, rotten food and fruits. It becomes a model organism for the study of diabetes due to fact that, about 75% of its genomes are functionally related to that of humans coupled with their small size and short generation time which makes it easy and inexpensive to culture in the laboratory (Millburn *et al*., 2016). *D. melanogaster* produce a large number of offspring within a short time (De-Campos *et al*., 2021). *D. melanogaster* possess powerful genetic contraptions that established its utilization as animal model for education and biomedical research (Millburn *et al*., 2016)

*Momordica charantia* is known as bitter melon or Garahuni in Hausa, Ejirin in Yoruba, is a plant that belongs to the order Cucurbitales and family *Cucurbitaceae. M. charantia* is widely distributed in tropical and subtropical regions of the world mainly in Asia where it is traditionally used as a medicinal plant and consumed as food (Villarreal-La *et al*., 2020). Reproductive fitness is the individual’s ability to fit in to the environment, pass on gene and produce more offspring to it subsequent generations (Singh and Rajender, 2022). Nutrition through consuming inappropriate resources depicts a challenge for organisms to reproduce due to high variation in the nutritional quality, which can result severe health performance and fitness (Ruendenauer *et al*., 2020).

## MATERIALS AND METHODS

### Study Area and Research Location

The study was conducted at the Drosophila and Neurogenetics Laboratory, Venom and Natural Toxins Research Centre (VANTRC), Ahmadu Bello University, Zaria (Latitude: 11º 8’ 18.938’ N and Longitude: 7º 38’ 43.052’ E), a suburb of Zaria in Kaduna State, Nigeria. Samaru, where the university is located, lies at Latitude: 11º 9’14.07” N. and Longitude: 7o 37’ 22.25” E.

### Collection of *M. charantia*

Fresh vegetative parts of *M. charantia* were collected from the wild in Gashua, Bade Local Government area, Yobe State (Latitude: 12° 51’ 41.33” N and Longitude: 11° 2’ 45.82” E) early in the morning (7:00 am) (Plate I). The samples were placed in appropriately labelled polythene bags, and transported to the Herbarium of the Department of Botany, A.B.U. Zaria for authentication using reference keys. The plant was assigned a voucher number: *ABU01597*.

**Plate I:**
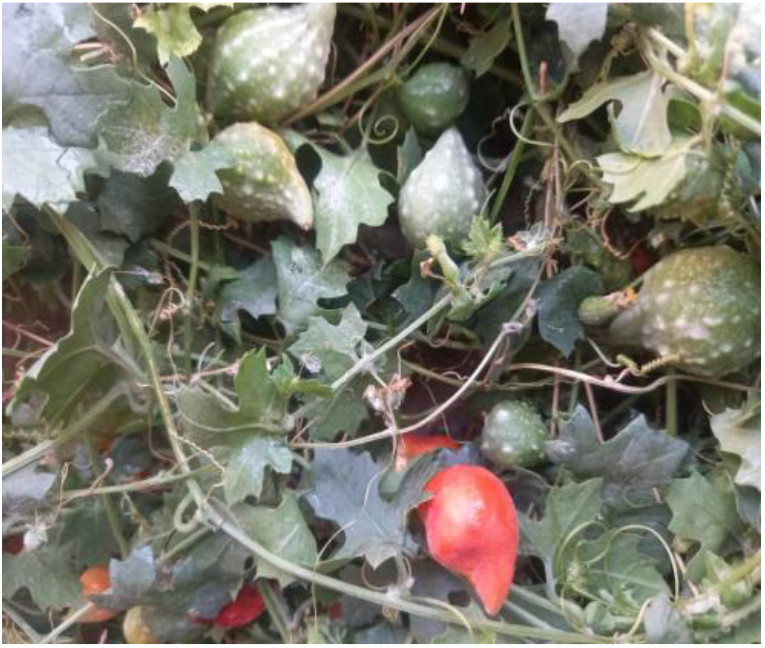
*Mormodica charantia*

### Ethanol Extraction from Leaves of *M. charantia*

Leaves of *M. charantia* were washed and shade-dried at room temperature for one week. The dried leaves were ground into powder using mortar and pestle. The powder was subjected to extraction using ethanol in a 1:10 ratio through three repeated successive macerations for 48 hours. The combined extracts were filtered, and the filtrate was evaporated under a vacuum at 40 °C using a rotary evaporator (Memmert, Schwabach, Germany). The resulting crude extract was stored in the refrigerator (N’Goran *et al*. (2019).

### Phytochemical Screening

Qualitative phytochemical screening of the ethanolic leaf extract was carried out according to the procedures stated by Vishnu *et al*. (2019)

### Gas Chromatography Mass spectrophotometry Analysis of the *M. charantia*

Portion of the ethanolic extract was dissolved in a 1:10 v/v solvent ratio, filtered filtered using a 0.45 μm nylon microfilter, and transferred to a sample vial (2 μl). The sample was injected into a GC-MS system (Model GC7890B 5977A MSD, Agilent Technologies, USA) at an injection port of 250 ºC and a split ratio of 5:1. The GC oven temperature program began at 110 ºC (2 mins hold), a increased at 10 ºC/min to 200 ºC (0 min hold) then at 5 ºC/min to 280 ºC (9 min hold). The sample was volatilized and separated into various components of ions of mass to charge ratio m/z, then transferred to the mass selective detector. The resulting mass spectra were analyzed using a standard reference library for compound identification.

### Meal Preparation

Cornmeal was prepared according to the method described by Markow and O’Gardy (2006). A total of 850 ml of distilled water was measured 700 ml was transferred into a pot and brought to a boil, while 150 ml was used to dissolve 50g of corn flour. Separately, 10g of yeast was dissolved in a portion of the boiled water, and 8g of Agar-Agar was added into the boiling water, stirring periodically to avoid lumps. After cooking for 10 minutes, the dissolved corn flour solution was added, cooked for another 10 minutes, followed by the yeast solution left to cook for another 15 minutes. The mixture was allowed to cool slightly before adding 1g of Nipagin dissolved in 5 ml ethanol.

### Induction of Type 2 diabetes

Type 2 diabetes was induced in strains of *D. melanogaster* by incorporating 2.5 g sucrose per 10 g of standard diet; all other ingredients of the standard fly-food were kept constant. After 10 days, the flies were assessed for diabetic symptoms such as reduced egg laying, smaller larval and adult sizes, and decreased locomotor activity (Omale *et al*., 2021).

### Filial Generation Assay

Three newly enclosed strains of *D. melanogaster* adult male and nine adult female flies were placed in vials containing respective diets. This generation of flies were recorded as parental (P) generation. Parental flies were removed after 48hrs to ensure synchronous larval growth. The larvae were allowed to puparite, rear and emerge as F1 flies. The F1 generation flies were counted and recorded. From the F1 generation, three male and nine female flies were selected randomly and transferred into fresh diets, to give rise to second filial generation (F2). This procedure was repeated to the 3rd filial generation (F3) (Chattophadhyay *et al*., 2015).

### Fecundity assay

A newly enclosed female was transferred in an egg laying chamber daily. The number of eggs were counted daily for 10 days. Five replicates plates were set up for each group (Chattophadhyay *et al*., 2015)

### Rapid Iterative Negative Geotaxis (RING) Assay

The locomotor performance of both the diabetic and treated flies were determined using the negative geotaxis method as previously described (Adedara *et al*., 2016). After 10 days of treatment ten flies were transferred into polystyrene vials, and allowed to recover. The vials were placed in a RING apparatus and acclimated for 15–20 minutes. The apparatus was tapped three times to knock flies down, and their climbing activity over 10 seconds was recorded. This was repeated five times at one-minute intervals. Images were scored for mean climbing height per vial.

### Data Analyses

The data obtained were analyzed using GraphPad Prism version 10 and excel, The results were expressed as mean ± standard error mean (SEM). The fecundity and filial generation were analyzed using two-way analysis of variance (ANOVA). *Tukey’s post-hoc* test was used to compare the level of significance across the groups. Geotaxis and larval crawling were done using One way ANOVA followed by *Tukeys post hoc* test, to compare the level of significance across the groups with P<0.05 considered statistical significant.

## RESULTS

### Phytochemical Composition of *M. charantia*

Phytochemical screening revealed the presence of alkaloids, cardiac glycosides, carbohydrates, tannins, saponins, flavonoids, terpenoids, steroids, and phenolic compounds; anthraquinones were absent (Table 1). GC-MS analysis identified 22 bioactive compounds in the ethanolic leaf extract, with 7, 11-Hexadecadienal having the highest (83.02%) Peak Area and cis-Vaccenic acid had the least (0.01%) Peak Area (Table 2).

**Table 1:** Secondary Metabolites of *M. charantia*.

| Phytochemical | Inferences |
| --- | --- |
| Alkaloids | + |
| Cardiac Glycosides | + |
| Saponins | + |
| Phenolic compounds | + |
| Tannins | + |
| Steroids | + |
| Carbohydrates | + |
| Flavonoids | + |
| Terpenoids | + |
| Anthraquinones | - |

**KEY:** + =Presence, - =Absence

**Table 2:** Primary Metabolites of *Mormodica charantia* Ethanolic Leaf Extract from GC-MS profiling.

| SN | Retention time (Minute) | Bioactive Compounds | % Peak Area |
| --- | --- | --- | --- |
| 1 | 10.45 | Hexadecanoic acid, ethyl ester | 0.49 |
| 2 | 11.47 | Oxirane, [(dodecyloxy)methyl]- | 0.11 |
| 3 | 11.64 | 2-Piperidinone, N-[4-bromo-n-butyl]- | 0.18 |
| 4 | 12.10 | 9,12-Octadecadienal | 0.62 |
| 5 | 12.42 | Ethyl 13-methyl-tetradecanoate | 0.42 |
| 6 | 16.41 | Bis(3-methylbutan-2-yl) phthalate | 0.30 |
| 7 | 19.30 | Oxalic acid, cyclobutyl hexadecyl ester | 0.13 |
| 8 | 20.20 | Squalene | 1.10 |
| 9 | 23.64 | 4-Fluoro-1-methyl-5-carboxylic acid, ethyl(ester) | 0.14 |
| 10 | 24.00 | trans-Traumatic acid | 0.21 |
| 11 | 24.10 | Oleic Acid | 0.18 |
| 12 | 24.38 | Oleic Acid | 0.14 |
| 13 | 25.25 | Oleic Acid | 2.04 |
| 14 | 25.62 | Disparlure | 0.49 |
| 15 | 26.47 | Oleic Acid | 2.75 |
| 16 | 26.84 | Z-8-Pentadecen-1-ol acetate | 0.98 |
| 17 | 27.17 | Cyclohexanol, 2-(2-propynyloxy)-, trans- | 1.68 |
| 18 | 27.27 | Z-17-Nonadecen-1-ol acetate | 0.36 |
| 19 | 27.69 | 1-Methoxymethoxy-oct-2-yne | 2.42 |
| 20 | 27.79 | 2-Methyl-Z,Z-3,13-octadecadienol | 0.72 |
| 21 | 28.07 | Oleic Acid | 1.37 |
| 22 | 30.30 | 7,11-Hexadecadienal | 83.02 |
| 23 | 32.93 | Undec-10-ynoic acid, tetradecyl ester | 0.10 |
| 24 | 32.99 | cis-Vaccenic acid | 0.01 |
| 25 | 33.42 | 9-Oxabicyclo[6.1.0]nonane, cis- | 0.09 |
| 26 | 33.85 | Oleic Acid | 0.06 |

### Filial generation of Harwich strain diabetic *D. melanogaster* fed with *M. charantia*

The result in Figure 1. The offspring numbers decreases from F1 to F3 in most groups; in the Negative control, the offspring count remains relatively stable across generations. A significant decreased in the Positive Control across the generation from F1 to F3. A 16 mg Met and 100 mg MC (26.47, 27.83) these groups show improvement over the positive control, particularly in F2 and F3. At 150 mg and 200 mg MC (21.27, 22.32), these groups do not show a strong recovery across generations, implying that higher doses could have diminishing effects over time.

**Figure 1.**
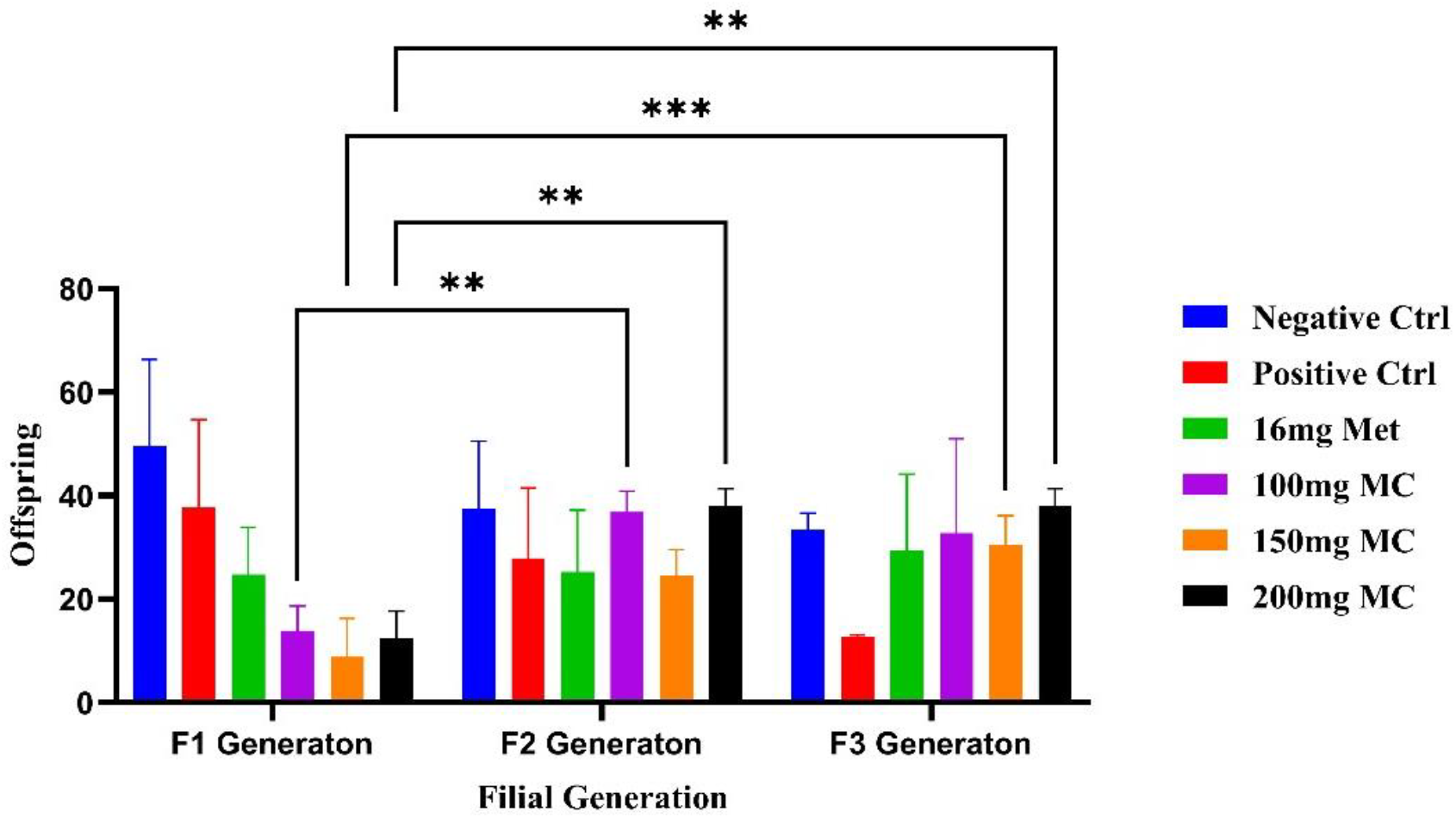
Filial generation of Harwich strain diabetic *D. melanogaster* fed with *M. charantia*.

### Filial generation of indigenous strain diabetic *D. melanogaster* fed with *M. charantia*

With F1 showing the most variability, while F2 and F3 display some recovery in certain treatment groups.Negative control offspring numbers remain relatively stable across generations. A decreased observed Positive control, particularly in F2, suggesting a persistent negative effect over generations. At 16 mg Met and 100 mg MC (26.63, 23.1), these treatments show a significant recovery from the negative effects observed in the positive control, especially in F2 and F3. At 150 mg and 200 mg (18.33, 15.63) MC These groups show limited or inconsistent recovery across generations, (shown in figure 2).

**Figure 2.**
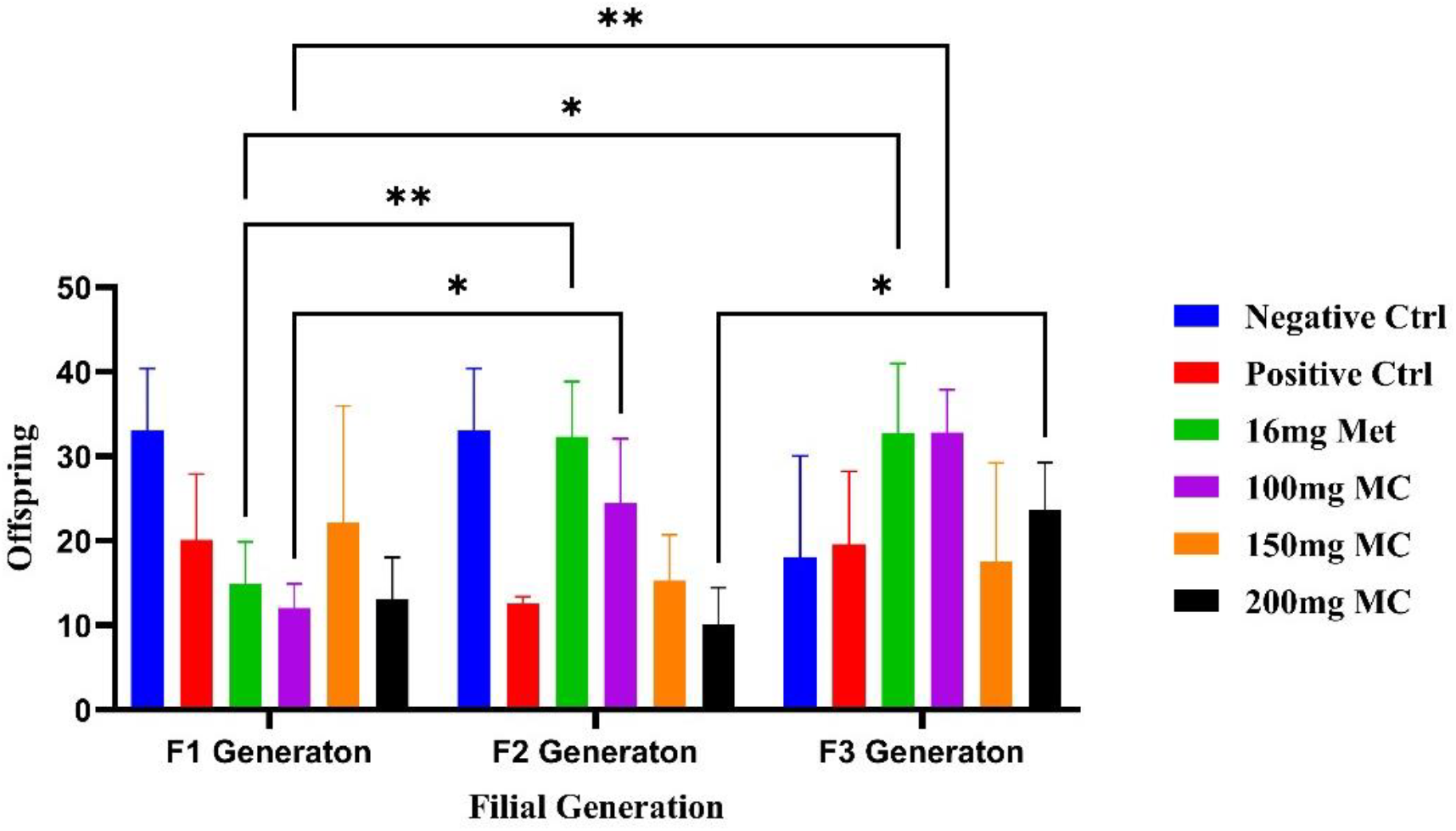
Filial Generation Output of Indigenous Strain of *D. melanogaster* Fed with *M. Charantia*.

### Fecundity Assay of Harwich strain diabetic *D. melanogaster* fed with *M. charantia*

In figure 3, the result shows that high sucrose (induced diabetes) did affect the flies ability to lay eggs, which was mostly improved when flies were co-exposed to 200 mg *M. charantia*. The highest number of eggs laid for most of the days was observed in 0.2 g *M. charantia* (with mean values of 7.8, 8.6, 9.8, 7.8, and 6.6 at days 2, 3, 4, 5, and 6). While the least was observed in untreated diabetic flies (positive control) with an average number of eggs (mean values 0.4, 0.2, 0.2, 0.4, and 0.2 at days 1, 2, 4, 5, and 6).

**Figure 3.**
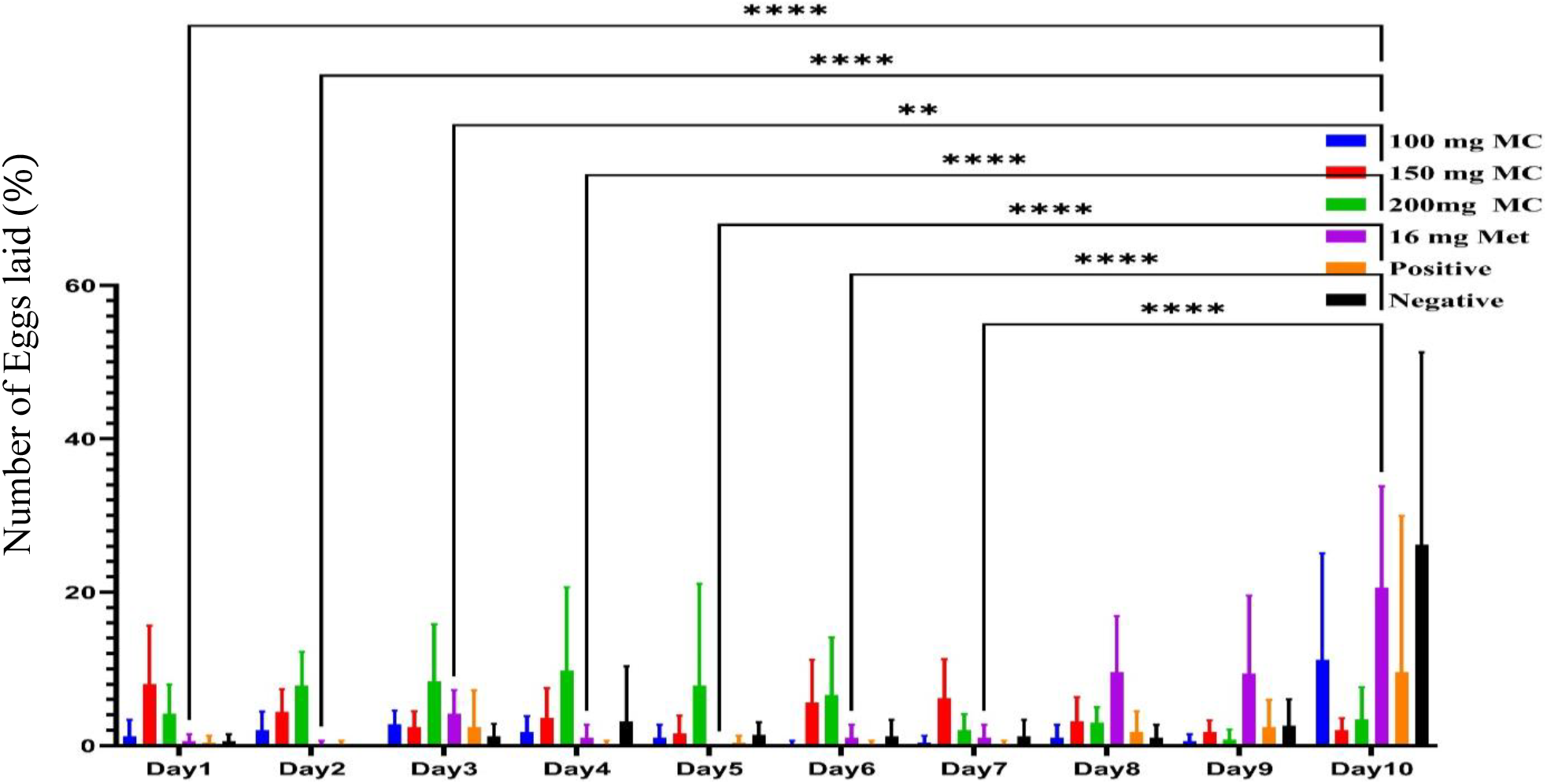
Fecundity Assay of Harwich strain diabetic *D. melanogaster* fed with *M. charantia*.

### Fecundity Assay of Indigenous strain diabetic *D. melanogaster* fed with *M. charantia*

The fecundity was presented in figure 4 below; the results showed that there is a significant (P<0.05) increase in eggs laid, flies fed with high sucrose, and treated with *the M. charantia* diet in three concentrations (100 mg, 150 mg, and 200 mg) in five replicates. *M. charantia* at a dose of 150 mg *M. charantia was* seen to have the highest number of eggs laid on days 1, 3, 4, and 6 with an average mean (5), while the least eggs laid was observed in diabetic (untreated) with an average mean value (0.5).

**Figure 4.**
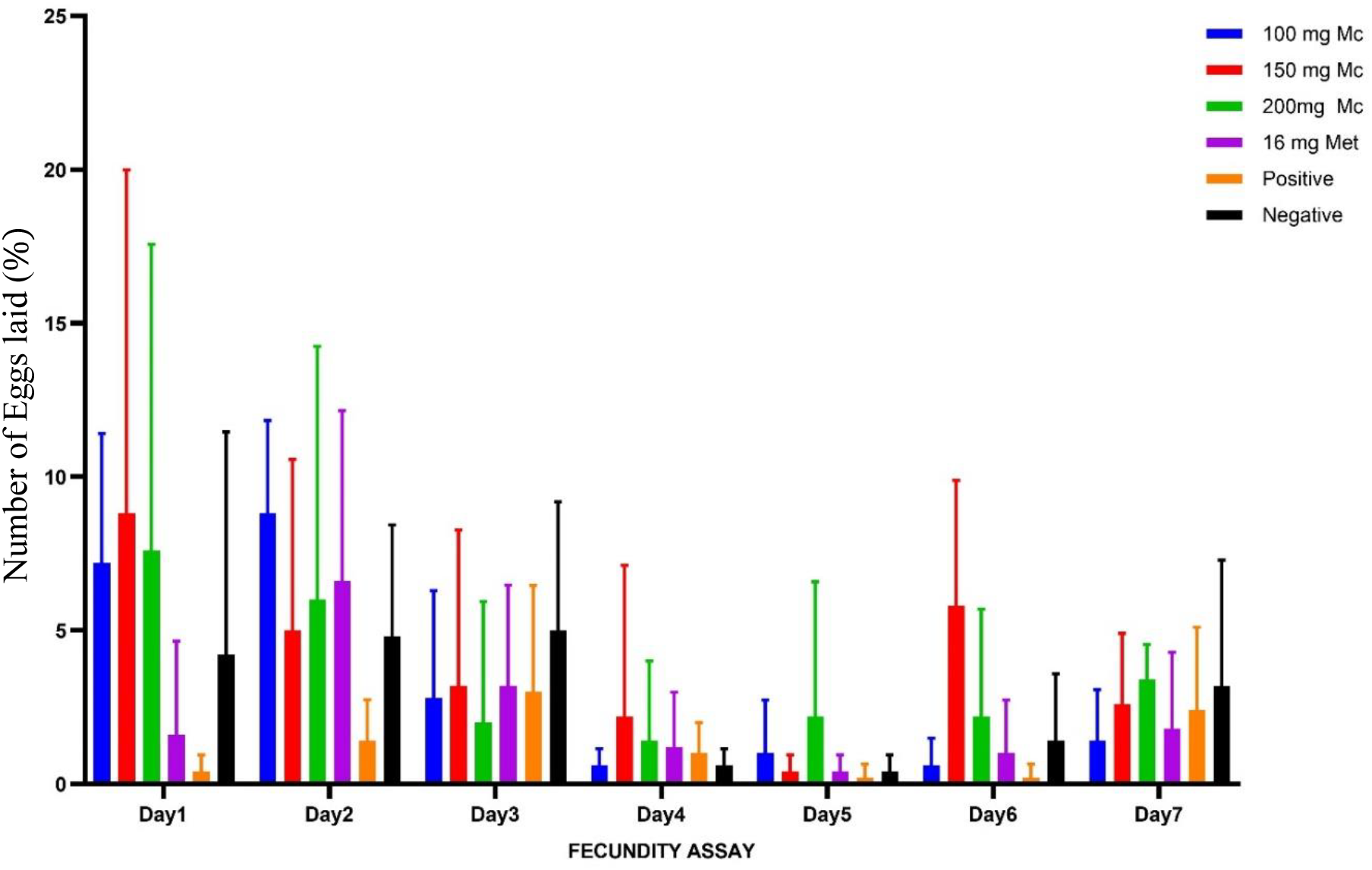
Fecundity Assay of Indigenous strain of *D. melanogaster* fed with *M. charantia*.

### Locomotor Performance

In the Harwich strain, *M. charantia* reduced locomotor performance in a dose-dependent manner, with the highest dose (200 mg) causing the greatest impairment (2.6) In contrast, in the Ngd3 strain, 150 mg *M. charantia* significantly improved locomotor activity (8.6) compared to the diabetic control (P < 0.01), whereas the 100 mg dose (4.2) showed reduced efficacy. Comparison between strains indicated that Ngd3 exhibited significantly better climbing accuracy than Harwich at 150 mg and 200 mg *M. charantia* (P < 0.001) (Figure 5).

**Figure 5.**
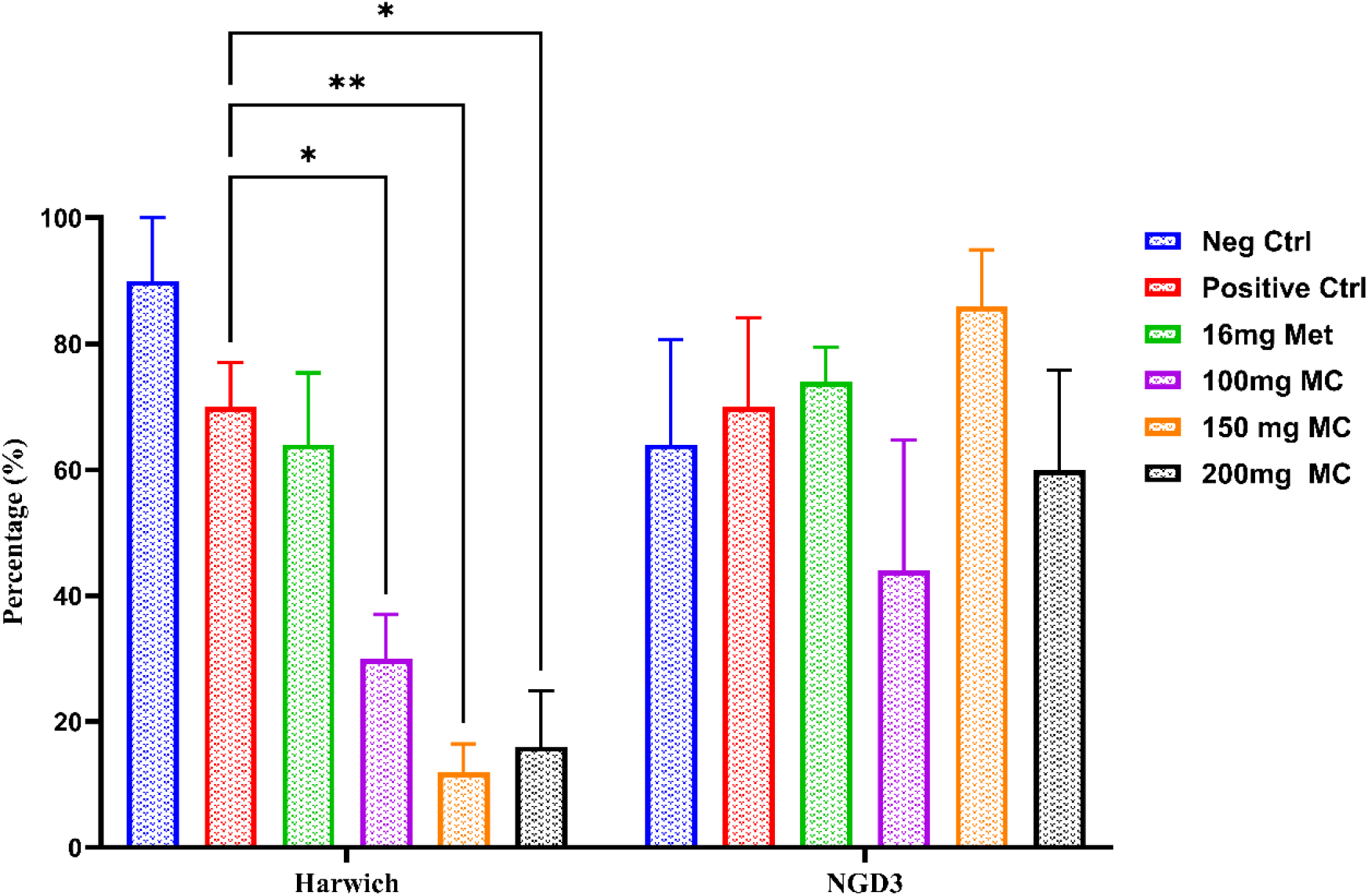
Geotaxis Assay of Harwich and Ngd3 Strains. n=10, mean ± SEM, two-way ANOVA, *Tukey post hoc* test, *=*p*<0.05, and **=*p*<0.001

## DISCUSSION

This study explored the anti-diabetic potential of *M. charantia* L. ethanolic leaf extract using *D. melanogaster* as a model for Type 2 Diabetes Mellitus (T2DM). Preliminary qualitative phytochemical analysis confirmed the presence of alkaloids, cardiac glycosides, saponins, phenolic compounds, tannins, steroids, carbohydrates, flavonoids, terpenoids in the ethanolic extract of *M. charantia* leaves. The phytochemicals profile obtained in this study is similar to the reports of Ayeni, *et al*. (2015) and Oyelere *et al*., (2022), but contrary to the reports of Suman and Jaya (2017), who reported the absence of terpenoids, steroids, and phenolic compounds in *M. charantia*. The differences in compounds may be attributed to differences in plant part, geographical origin, and extraction methodology. The presence of phenolics and flavonoids is particularly worth highlighting, as these compounds possess potent antioxidant and insulin-sensitizing properties. Such bioactivities are critical in ameliorating hyperglycemia and oxidative stress; two major hallmarks of T2DM. Additionally, these secondary metabolites have been associated with enhanced pancreatic β-cell function and improved glucose metabolism (Suman and Jaya, 2017), suggesting a potential mechanistic basis for the observed therapeutic effects.

The identification of bioactive compounds in *M. charantia* lays the foundation for new avenues in drug discovery and development, particularly due to the presence of significant secondary metabolites (Srinivasulu *et al*., 2017). Among these, saponins from *M. charantia* have been shown to activate AMP-activated protein kinase (AMPK) thereby inhibiting gluconeogenesis in the liver of diabetic mice (Wang *et al*., 2019). Saponins exert their anti-hyperglycemic effects through multiple mechanisms, including of protection of pancreas β-cells, stimulation of insulin secretion and improvement of ameliorating insulin sensitivity (Marrelli *et al*., 2016; El barky *et al*., 2017). Flavonoids and other polyphenols have also demonstrated blood glucose-lowering properties by enhancing GLUT-2 in pancreatic β-cells (Belayneh *et al*., 2019; Melaku and Getnet, 2020). Promoting secretion (Khalid *et al*., 2019), and increasing the expression and translocation of GLUT-4 transporters (Moradi *et al*., 2018).

Furthermore, flavonoids are known to slow gastric emptying and inhibit carbohydrate-digesting enzymes such as α-glycosidase and α-amylase (Kifle *et al*., 2020; Gebremeskel *et al*., 2020). Tannins and phenolic compounds may contribute to glucose lowering effects by stimulateing insulin secretion or mimicking insulin activity, reducing carbohydrate absorption through enzyme inhibition, and protecting β-cells via antioxidant mechanisms (Melaku and Getnet, 2020). In addition, alkaloids have garnered increasing interest for their potential anti-diabetesic properties, particularly through the inhibition of α-glucosidase and α-amylase (Kumar *et al*., 2019, Bhaskaran *et al*., 2019).

Gas Cchromatography-Mass Spectrophotometry analysis revealed 26 bioactive constituents, including 2-piperidinone, squalene, phytol, hexadecanoic acid ethyl ester, and bis(3-methylbutan-2-yl) phthalate. These compounds are known for their antioxidant, anti-inflammatory, hypolipidemic, and glucose-modulating properties (Rani *et al*. 2009; Ponnamma and Manjunath 2012; Gomathi *et al*., 2015). Esters are important organic compounds that increases number of commercial applications (Foresti *et al*., 2005). These compounds are largely used in fragrances, cosmetics detergents, flavours and pharmaceuticals. Due to the presence of above mentioned compounds in the whole plant ethanolic extract of *M. charantia* may be used in various pharmaceutical and industrial applications.

The result of this study shows the effect of *M. charantia* on the output filial generation of Harwich diabetic *D. melanogaster*. The flies co-treated with 200 mg *M. charantia* produced the most across three generations, which is similar to previous findings and suggested that MC extract enhanced the number of offspring in *D. melanogaster* reproductive fitness (Fan *et al*., 2019). While those treated with metformin produced the lowest support, the alternative treatments for diabetes, such as the use of MC, could provide supplementary benefits (Brouwer *et al*., 2018). Similarly, Kumar *et al*. (2018) observed that *M. charantia* supplementation improved reproductive success in *Drosophila melanogaster*.

This study explored the pharmacodynamics of *M. charantia* on diabetic Harwich strain *D. melanogaster* reproductive fitness. The flies subjected to the highest concentration of 200 mg of MC had the highest number of eggs laid, while the positive group had the least number of eggs laid, which suggests that diabetes affects the fecundity ability of the flies. The increased egg-laying ability observed in flies treated with higher concentrations of MC aligns with findings from previous studies and could suggest the beneficial effects of *M. charantia* in enhancing reproductive parameters in various models. For instance, studies have shown that the bitter melon extract positively influences reproductive functions by modulating insulin sensitivity and reducing oxidative stress, which are often exacerbated in diabetic conditions (Zhang *et al*., 2020). In contrast, the lower reproductive output in the metformin-treated group could indicate that while the metformin is effective in managing blood glucose levels, it may be the reproductive impairments linked to diabetes, thereby supporting the idea that alternative treatments like MC could provide supplementary benefits (Brouwer *et al*., 2018).

In the case of the filial generation output of the indigenous strain *D. melanogaster, the* number of offspring was also significantly reduced by high sucrose, which was rescued at a dose of 100 mg MC and 16 mg Met. This enhancement could be a result of the hypoglycemic effects of *M. charantia*, as previously reported by Xu *et a*l. (2022). The health-promoting properties of *M. charantia*, such as anti-fatality, hypoglycemia, and antimutagene, may contribute to its great potential effects as a medicinal plant (Jia *et al*., 2017). This study is in contrast to the finding of AC *et* al. (2011), who reported that 25 mg/100 g of *M. charantia* significantly improved the diestrous phase of Sprague-Dawley rats.

Fecundity was assessed over a 7-day period; the egg-laying ability of indigenous strain *D. melanogaster* was significantly reduced by the high sucrose (non-treated diabetic flies), which was mostly improved by 150 mg MC. This improvement can be attributed to the antidiabetic properties of *M. charantia*, as previously reported (Akaneme, 2008). The phytochemical constituents of *M. charantia*, such as charantin, vicine, and polypeptide-p, may contribute to its hypoglycemic effects (Oyelere *et al*., 2022).

This finding is similar to the study conducted by Hussain *et al*. (2022), who reported that 500 mg/kg of *M. charantia* significantly enhanced ovulation in Swiss albino adult female rats.

Behavioural assays provided additional insights into the functional recovery of diabetic flies. The neuromuscular test (climbing assay) revealed sensory-motor deficits under diabetic conditions, more pronounced in the Harwich strain. The Harwich strain performed better in the absence of any treatment but did not show same level of improvement with *M. charantia*. Treatment with 100 mg *M. charantia* restored performance, indicating neuroprotective effects, while higher doses (150 – 200 mg) produced diminished or adverse outcomes, possibly due to dose-related toxicity or interference with normal metabolic processes. This dose-dependent decline is consistent with earlier studies that reported adverse effects of high doses of *M. charantia*, particularly in systems under metabolic stress, such as in hyperglycemic conditions (Joseph and Jini, 2013). This biphasic response highlights a critical therapeutic window; low to moderate doses are beneficial, whereas higher doses may impair neuromuscular coordination or induce oxidative stress (Basch *et al*., 2003; Nerurkar *et al*., 2010). When compared to Metformin, the optimal does of the extract produced similar behavioural improvements, further emphasizing its potential to alleviate diabetic neuropathy and motor dysfunction.

The significant improvement in climbing abilities in the NgD3 strain treated with *M. charantia* suggests that the extract may offer cognitive or motor benefits in diabetic models at an optimal dose of 150 mg, potentially through mechanisms involving neuroprotection or enhanced glucose metabolism in the brain. Previous studies have reported that *M. charantia* possesses neuroprotective properties and can improve cognitive function in diabetic models (Kumar *et al*., 2010). Higher doses of *M. charantia* significantly improved accuracy in the Ngd3 strain, which may have a genetic predisposition to respond more favorably to such treatments. The findings of this study are consistent with the report of Omale *et al*. (2021), who reported the glucose-lowering effect of *Parinari curatellifolia* stem bark in HDS-induced flies.

## CONCLUSION

This study highlights the significant anti-diabetic potential of *M. charantia* in improving reproductive fitness and neuromuscular function in T2DM-induced *D. melanogaster*. The 100 mg dosage demonstrated the most effective results in mitigating hyperglycemia-induced neuromuscular dysfunction and enhancing reproductive parameters. These findings suggest that *M. charantia* could serve as a promising natural alternative for diabetes management. However, the dose-dependent effects observed indicate the necessity for further investigations into optimal therapeutic dosages. Studies could be focus on the specific pathways well as assessing the long-term impact of MC treatment on neuromuscular function and overall health in hyperglycemic models

